## Supplemental Figures for "Dynamics of mesoscale brain network during visual discrimination learning revealed by chronic, large-scale single-unit recording"

### Figure supplements

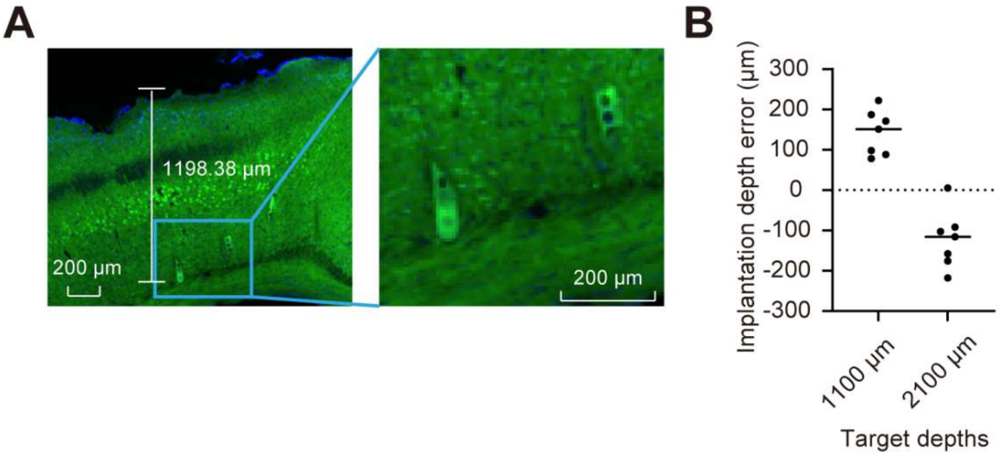


**Figure S1, related to Figure 1. Implantation depth accuracy test.** (A) Example section showing recovered array shanks in the cortex. The actual implantation depth was defined according to the location of the guiding hole at the tip of the shanks. (B) Errors in implantation depth, as the difference between actual depth and original target depth (negative values represent shallower depth). A slight tendency for deeper errors in cortical implantation (1100 μm) and shallower errors in subcortical implantation was observed.

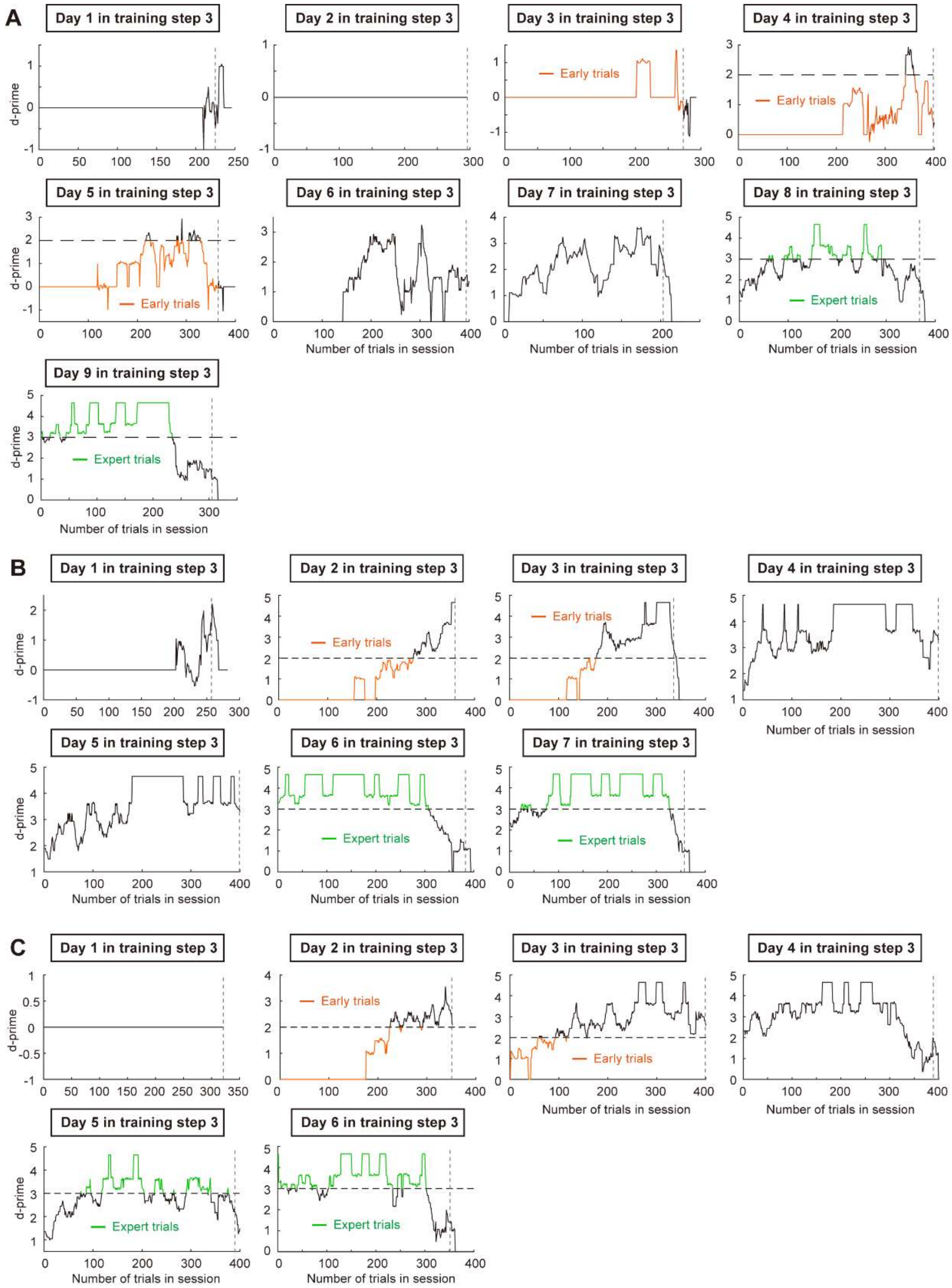


**Figure S2, related to Figure 1. Behavioral performance in daily sessions.** (A-C) D-prime curves for the three mice in the learning group. Each data point represents the behavioral discriminability [17] calculated from the 10 trials before and 10 trials after the corresponding trial. Although the d-prime overall increased with training, the task performance showed nonnegligible fluctuations in each session. The trials after the last licking (indicated by vertical dashed lines) in each session were excluded. Colored segments mark the data used in subsequent analyses. For data at the early training stage, trials in early sessions with d-prime < 2 (orange lines) were used, and for expert stage data, trials in late sessions with d-prime > 3 (green lines) were used.

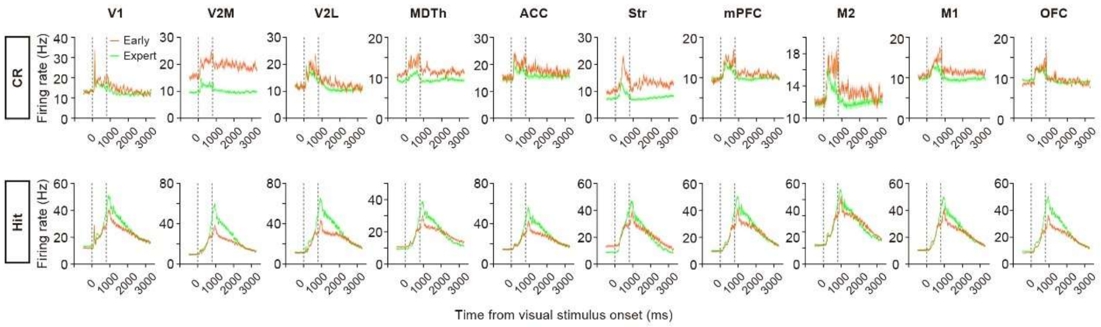


**Figure S3, related to Figure 2. Average firing rate from the bootstrap-resampled datasets.** The mean traces of all bootstrap-resampled datasets (n = 500 datasets) were plotted, and were highly similar to original data. Firing rate traces were aligned to the visual stimulus onset. Shading, SEM.

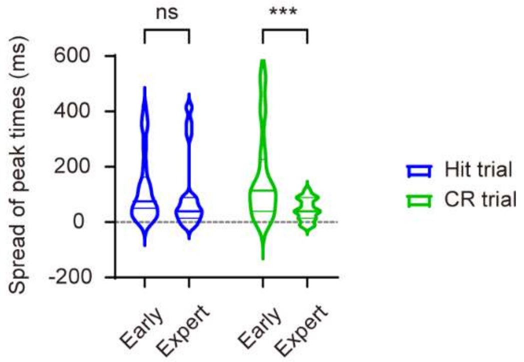


**Figure S4, related to Figure 2. Spread of peak activation times across brain regions.** Violin plot showing the distribution of differences between peak activation times of each region pairs at different learning stages. Thick lines in the boxes indicate medians, thin lines in the boxes indicate quartiles. ***$p < 0.001$, t-test with Sidak correction.

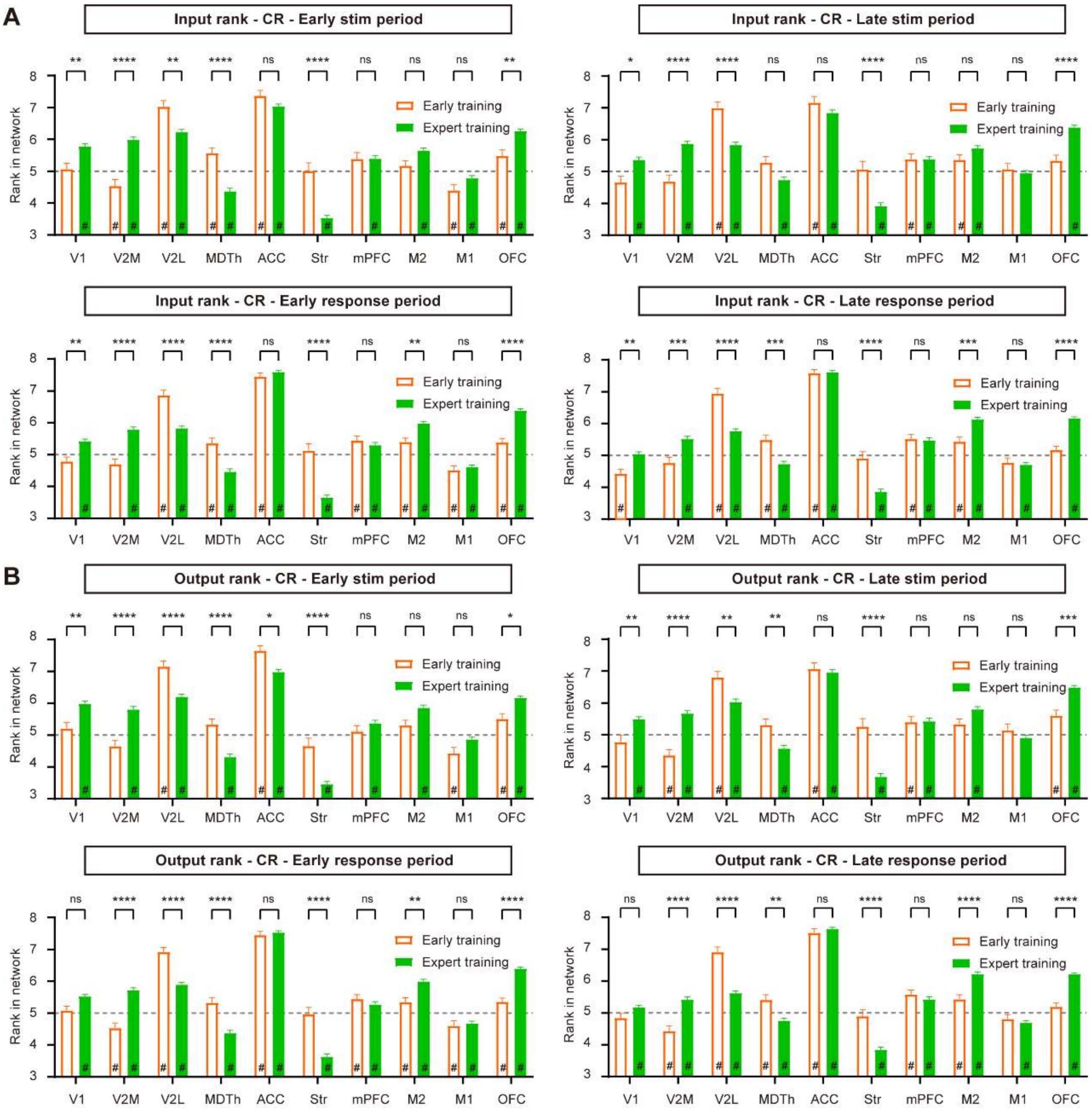


**Figure S5, related to Figure 4. Ranking dynamics in CR trials during learning.** (A) Average input rank of each brain region in the early stimulus period (0-400 ms after stimulus onset), late stimulus period (400–800 ms after stimulus onset), early response period (800–1800 ms after stimulus onset), and late response period (1800–2800 ms after stimulus onset) of early and expert CR trials. (B) Same as A but for output ranks. *$p < 0.05$, **$p < 0.01$, ***$p < 0.001$, ****$p < 0.0001$, t-test with Sidak correction. # indicates the rank value significantly different ($p < 0.05$, t-test) from 5 (average level of random data, indicated by dashed lines). n = 118 early CR trials from 7 sessions of 3 mice, and 610 expert CR trials from 6 sessions of same mice. Error bars, SEM.

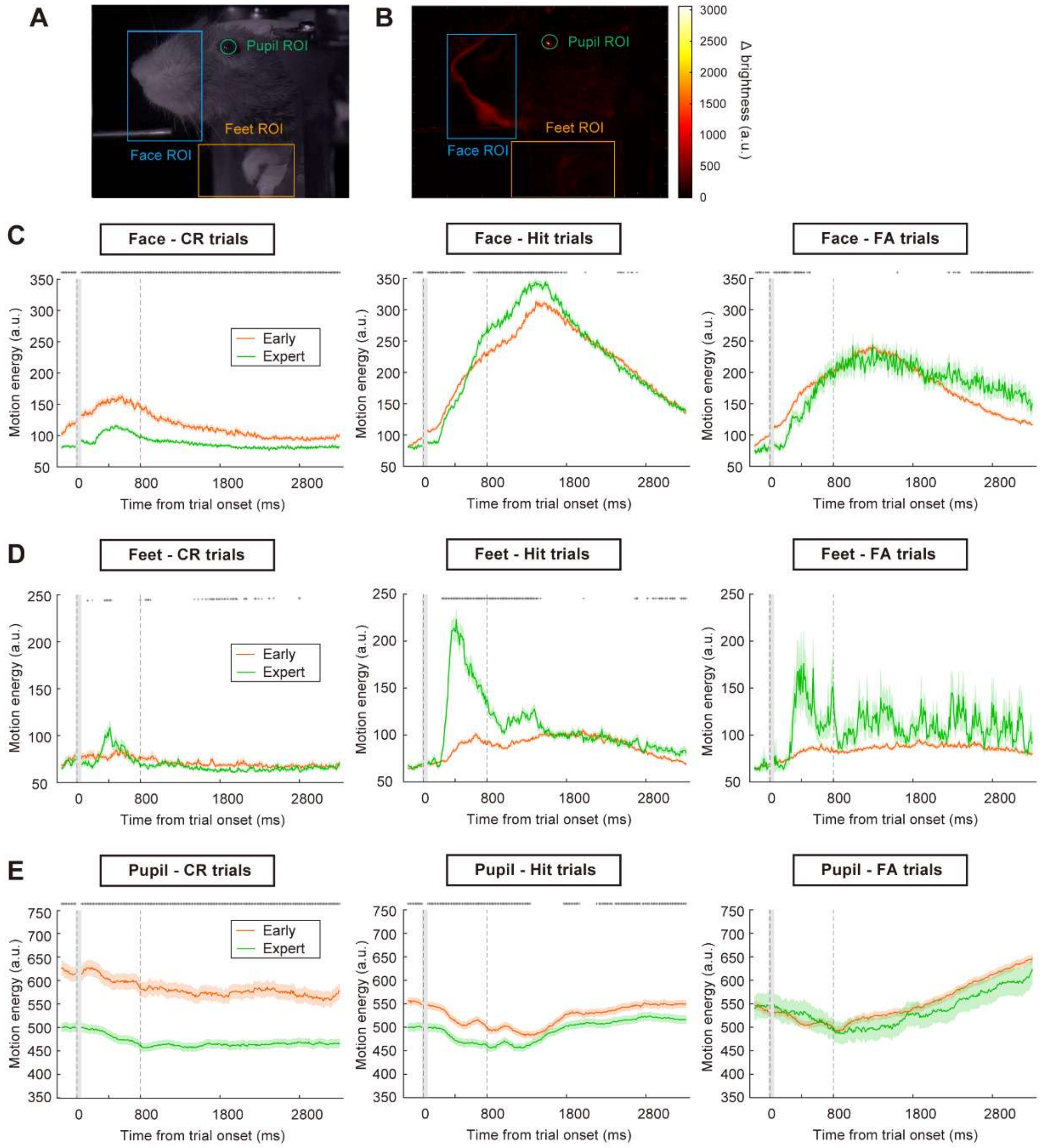


**Figure S6, related to Figure 4. Motion energy in CR trials during learning.** (A) example video frame showing regions of interest (ROIs) used in oral-facial movement analysis to measure motion energy [21]. (B) Example showing the absolute value of brightness difference between consecutive frames. (C-E) Average motion energy of mouse oral-facial movements in CR, Hit and FA trials. n = 466, 1422, 1184 early CR, Hit, FA trials and 730, 774, 92 expert CR, Hit, FA trials from 3 mice. *, $p < 0.05$, t-test with Sidak correction. Error bars, SEM.

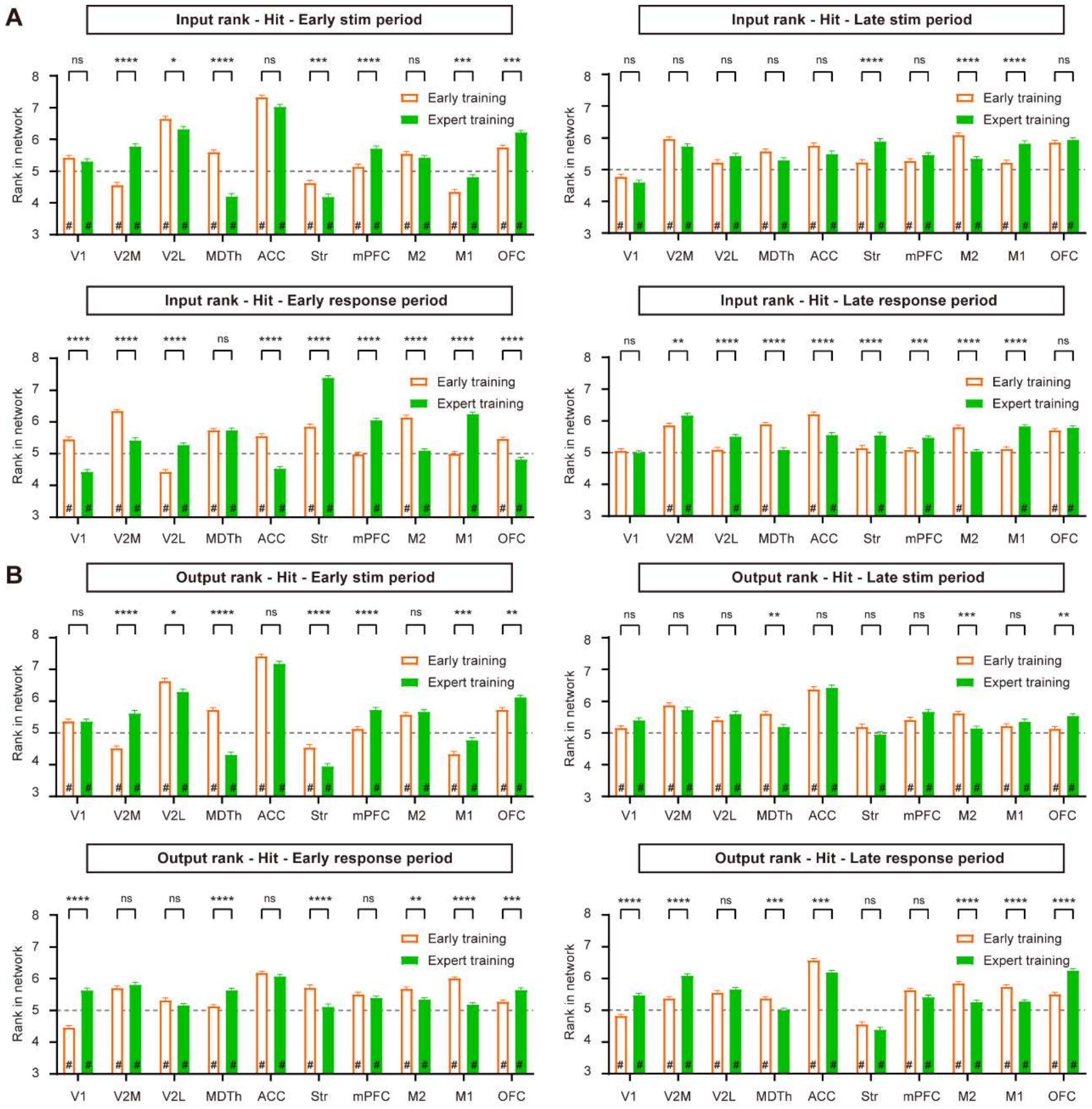


**Figure S7, related to Figure 5. Ranking dynamics in Hit trials during learning.** (A) Average input rank of each brain region in the early stimulus period (0–400 ms after stimulus onset), late stimulus period (400–800 ms after stimulus onset), early response period (800–1800 ms after stimulus onset), and late response period (1800–2800 ms after stimulus onset) of early and expert Hit trials. (B) Same as A but for output ranks. *$p < 0.05$, **$p < 0.01$, ***$p < 0.001$, ****$p < 0.0001$, t-test with Sidak correction. # indicates the rank value significantly different ($p < 0.05$, t-test) from 5 (average level of random data, indicated by dashed lines). n = 828 early Hit trials from 7 sessions of 3 mice, and 677 expert Hit trials from 6 sessions of 3 mice. Error bars, SEM.

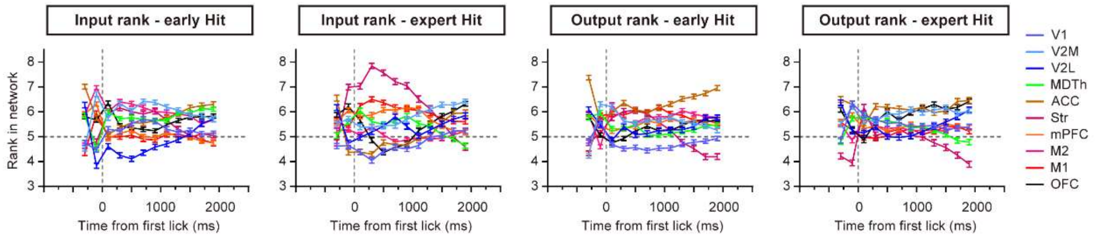


**Figure S8, related to Figure 5. Ranking dynamics in Hit trials during learning, with spike time aligned to the first lick of each trial.** Input/output ranking dynamics in early and expert Hit trials, with spike time aligned to the first lick of each trial and the cross-correlation analyses recalculated. Horizontal dashed lines indicate the average level of random data.

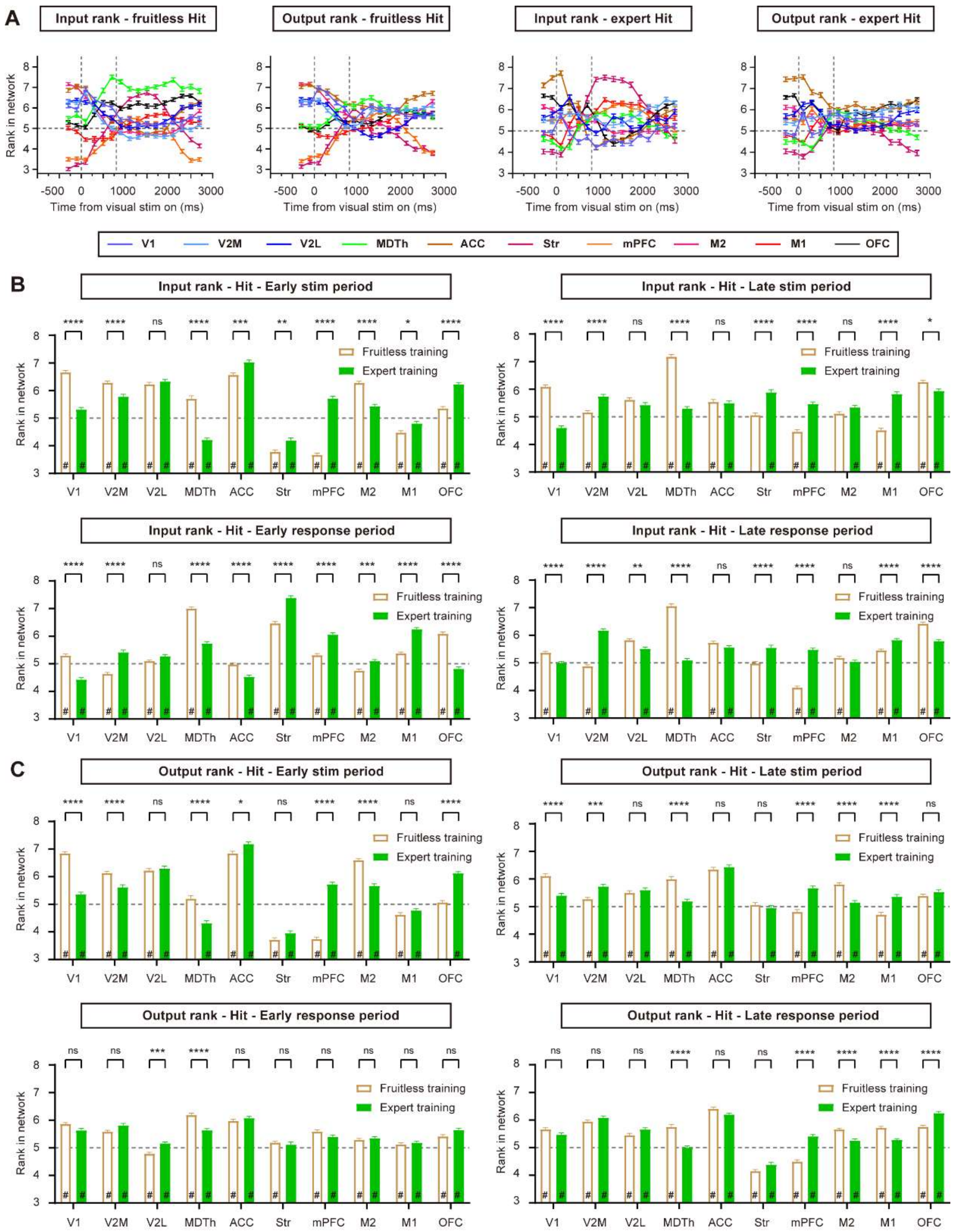


**Figure S9, related to Figure 5. Ranking dynamics in fruitless-learning Hit trials compared to expert Hit trials.** (A) Input/output ranking dynamics in the fruitless-learning Hit trials and expert Hit trials. (B) Average input rank of each brain region in the early stimulus period (0–400 ms after stimulus onset), late stimulus period (400–800 ms after stimulus onset), early response period (800–1800 ms after stimulus onset), and late response period (1800–2800 ms after stimulus onset) of the fruitless-learning and expert Hit trials. (C) Same as B but for output ranks. *p < 0.05, **p < 0.01, ***p < 0.001, ****p < 0.0001, t-test with Sidak correction. # indicates the rank value is significantly different (p < 0.05, t-test) from 5 (average level of random data, indicated by dashed lines). n = 844 Hit trials from 8 sessions of 2 fruitless-learning group mice, and 677 expert Hit trials from 6 sessions of 3 normal-learning group mice. Error bars, SEM.

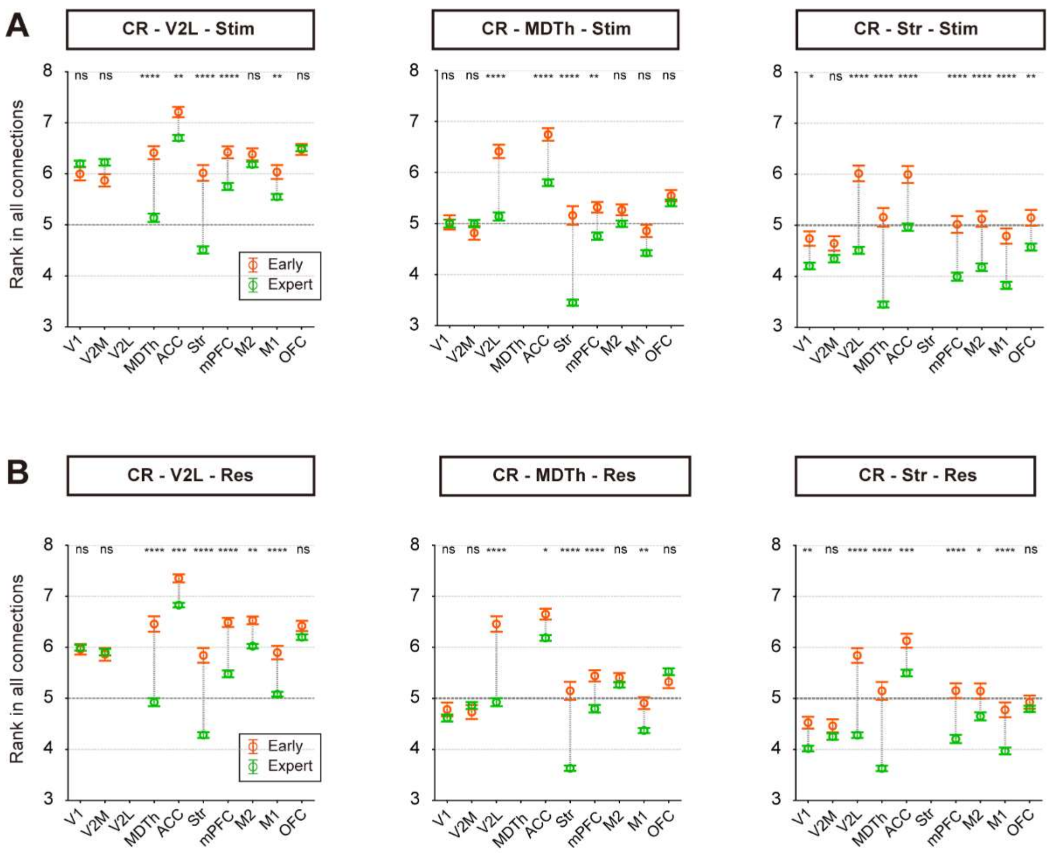


**Figure S10, related to Figure 6**. **Rank decrease in CR trials was attributed to reduced input/output rank from/to other regions.** (A) Average input rank changes for the three regions (V2L, MDTh, and striatum) that showed decreased rank values in CR trials during visual learning. *$p < 0.05$, **$p < 0.01$, ***$p < 0.001$, ****$p < 0.0001$, t-test with Sidak correction. (B) Same as A but for the response period of CR trials. (C-D) Same as A-B but for output ranks. n = 118 early CR trials from 7 sessions of 3 mice, and 610 expert CR trials from 6 sessions of 3 mice. Dashed lines at rank 5 indicate the average level of random data. Error bars, SEM. Stim, stimulus period. Res, response period.
